# Guanylyl cyclase signaling in AFD neurons regulates systemic stress resilience in *Caenorhabditis elegans*

**DOI:** 10.1101/2025.07.08.663649

**Authors:** Abhilasha Batra, Rati Sharma

## Abstract

Thermal fluctuations in the environment, particularly high temperatures, pose a major challenge to organisms and require robust mechanisms for survival under heat stress. Although the molecular basis of cellular heat shock responses is well understood, how thermosensory neurons contribute to systemic stress adaptation remains unclear. Here, using *Caenorhabditis elegans* as a model, we examine whether thermosensory receptor guanylyl cyclases (rGCs) in AFD neurons regulate organism-wide stress responses under noxious temperatures and how individual rGCs contribute to this coordination. Among the AFD-expressed rGCs, we identify *gcy-18* and *gcy-23* as key regulators of the physiological response to thermal stress, acting through modulation of canonical heat shock response (HSR) genes. Our findings indicate that rGC signaling is crucial for activation of heat shock chaperones and maintenance of proteostasis under high temperatures (35^°^C). Supporting this, quantitative analysis of HSP-16.2 revealed that rGC activity in AFD neurons modulates the HSR magnitude in distal tissues, such as the intestine. Together, our findings uncover an important role for thermosensory rGCs in maintaining cellular proteostasis through selective modulation of the HSR.

## INTRODUCTION

Temperature is an important environmental factor that influences the physiology, behavior and survival of organisms. In ectotherms like *Caenorhabditis elegans* (*C. elegans*), exposure to temperatures beyond the optimal range can trigger physiological challenges, including cellular stress, reduced lifespan and impaired development [1–3]. While these challenges are typically linked to the direct effects of high temperatures on protein stability and cellular functions, there is increasing evidence that organisms also possess mechanisms to modulate their response to heat-induced physiological stress. One well-known intracellular mechanism that enables organisms to cope with noxious temperature conditions is the heat shock response (HSR) pathway that leads to the expression of stress-inducible heat shock proteins (HSPs) [4, 5]. These proteins help provide cellular protection to the organism by repairing damaged proteins, thereby mitigating cellular stress and maintaining protein homeostasis even under extreme conditions [6–10].

In multicellular organisms with a nervous system, such as *C. elegans*, the sensory neuronal circuits also initiate many of the animals’ responses to environmental stimuli [11, 12], including chemicals, mechanical stretch, light and temperature. The neuronal system of *C. elegans* comprises of 60 ciliated sensory neurons, which serve as antennas for the detection of external cues [13]. Among these, the amphid finger-like AFD neurons, which terminate in sheath cells, are responsible for transmitting thermal information to interneurons for further processing. These AFD neurons are the major thermosensory neurons in *C. elegans*, that enable the worms to sense temperature changes and adjust their behavior and physiology to adapt to new environmental temperature. Under ambient temperature conditions (15-25^°^C), the thermosensory signaling by AFD neurons manifests in the thermotaxis behavior of worms, where animals cultivated at a certain temperature migrate toward that temperature on a thermal gradient [14]. AFD neurons mainly sense temperature changes through a molecular signaling cascade that is sensitive to thermal stimuli. Central to this cascade are three receptor guanylyl cyclases (rGCs), GCY-8, GCY-18 and GCY-23, which are expressed specifically in AFD and are responsible for the catalytic production of second messenger cyclic guanosine monophosphate (cGMP) in response to warming [15, 16]. The increased cGMP levels then activate the cyclic nucleotide-gated (CNG) channels, composed of TAX-4 and TAX-2 subunits. The opening of these channels lead to a calcium influx, which in turn activates the neuron [17–19]. The essential role of this neuronal signaling pathway is highlighted by various studies showing that animals lacking three of the AFD-specific rGCs or the TAX-4/TAX-2 channels exhibit defective thermotaxis and fail to show AFD response to warming [15, 16, 18, 20, 21].

The three rGCs, GCY-8, GCY-18 and GCY-23 expressed in AFD neurons function redundantly yet differentially to mediate thermo-responsiveness and to regulate thermotaxis. These findings support the idea that these rGCs serve as the primary temperature sensors in *C. elegans* [16]. The distinct roles of each GCY in modulating cGMP signaling within AFD neurons and their contributions to thermosensory responses at innocuous temperatures have been well characterized [15, 16]. However, *C. elegans* is capable of detecting not only innocuous temperatures but also noxiously high and low temperatures, as evidenced by the physiological and molecular changes elicited under such extreme conditions [22–24]. In fact, various studies in the past have shown that under noxious thermal conditions (*>*30^°^C), the protein folding environment and other cellular stress responses in *C. elegans* are regulated cell-nonautonomously by the thermosensory neuronal circuitry [25–28]. Furthermore, it has also been shown that worms carrying a mutation in the *gcy-8* gene exhibit mitigated HSR activation on exposure to heat [28], thus highlighting a link between the sensory signaling pathway and systemic stress responses. This cell-nonautonomous regulation of the HSR by the AFD thermosensory neurons has emerged as a key area of investigation, revealing intricate mechanisms by which sensory neurons influence the organism-wide response to thermal stress [29, 30].

Previous studies have investigated the role of rGCs in AFD neuronal signaling and their influence on chronic protein misfolding in *C. elegans* using loss-of-function mutations under short-term acute heat shock conditions (*i*.*e*., 34^°^C for 15 min) [24, 28, 29]. These studies have provided insights into how AFD neurons and rGCs contribute to thermosensory responses and stress signaling. However, the specific contributions of individual rGCs (GCY-8, GCY-18 and GCY-23) under prolonged exposure to noxious temperatures remain largely unexplored. In particular, it is unclear how each GCY, either independently or in combination modulates the physiological responses of *C. elegans* to sustained heat stress and influences the induction and regulation of HSR. Understanding how each of these rGCs functions under prolonged heat stress conditions is essential for elucidating how AFD neurons drive systemic stress adaptation in *C. elegans*.

To address this knowledge gap, we investigate the distinct role of three AFD-specific rGCs, GCY-8, GCY-18 and GCY-23, in regulating the physiology of *C. elegans* under prolonged noxious thermal stress. Specifically, we ask how sustained high temperature exposure affects the induction of HSR and the organism’s overall stress resilience, and how each GCY contributes to this process. To this end, we employ a systematic mutant framework approach comprising *gcy* single mutants, *gcy* double mutants that retain functionality in only one of the three *gcy* genes and a triple *gcy* mutant in which all three *gcy* genes are disrupted, thus effectively abolishing thermosensory reception. By comparing the stress phenotypes and HSR dynamics across these mutants, we aim to unravel the individual and combined contributions of these rGCs in mediating sensory-driven protection during thermal stress.

Through this approach, our study provides a comprehensive understanding of how AFD neurons via their associated rGCs coordinate systemic stress responses in *C. elegans*, particularly under prolonged noxious temperature conditions.

Here, we show that AFD expressed rGCs, GCY-8, GCY-18 and GCY-23, are not only involved in thermosensory signaling but also contribute significantly to heat stress adaptation in *C. elegans*. Through a combination of genetic and physiological analyses, we find that these rGCs in addition to their established roles in thermosensory transduction under optimal temperatures, are also essential for survival and recovery under prolonged noxious heat conditions. Notably, our results indicate that a subset of these rGCs is required for HSF-1 dependent activation of HSR, including the expression of key molecular chaperones such as *hsp-70* and *hsp-16*. This reveals a functional link between the receptors of the AFD thermosensory pathway and cellular proteostasis. Our findings then, in effect, establish that AFD neurons rely on their rGCs to transduce thermal cues into a cell-nonautonomous signal, which in turn drives protective transcriptional responses that are essential for heat stress adaptation in *C. elegans*.

## RESULTS

### AFD-rGCs are necessary for the survival of worms at elevated temperatures

To determine whether AFD-rGCs play a role in the thermotolerance ability of *C. elegans*, we examined the survival of wild-type and rGC mutant worms at 35^°^C over time (Fig. 1a-c) and quantified the TD50 (the time at which 50% of the population remains viable at 35^°^C) across different genotypes (Fig. 1d) [31]. As shown in Fig. 1a, we observed a significant reduced surviving fraction in *gcy-8 gcy-18 gcy-23* triple mutants of rGCs in comparison to the wild type, thus indicating a severe thermotolerance defect. While wild-type worms show a gradual decline in survival over the 10 h heat stress period, the *gcy* triple mutant population experiences a more rapid loss, indicating that AFD-rGCs collectively play a crucial role in maintaining heat resistance. To understand the contribution of individual rGCs to thermotolerance, we examined the survival rates of single mutants (*gcy-8, gcy-18* and *gcy-23*). As shown in Fig. 1b, the survival curves of single mutants largely overlap with that of wild type, with major reductions observed at certain time points in *gcy-18* and *gcy-23* mutants and only minor in *gcy-8*. Since none of the single mutants exhibit the pronounced decline in survival observed in the rGC triple mutant, the loss of thermotolerance cannot be solely attributable to any individual *gcy*, but likely arises from their combined activity. To further dissect the combinatorial effects of individual rGCs, we compared the survival of double mutants to their respective single mutant counterparts (Fig. 1c). We observed, in some cases, that the double mutants show a greater reduction in survival than single mutants, though the effect remained less severe than that observed in the triple mutant. This indicates that multiple rGCs contribute to the same thermotolerance pathway, with each playing a distinct yet overlapping role.

**Figure 1.**
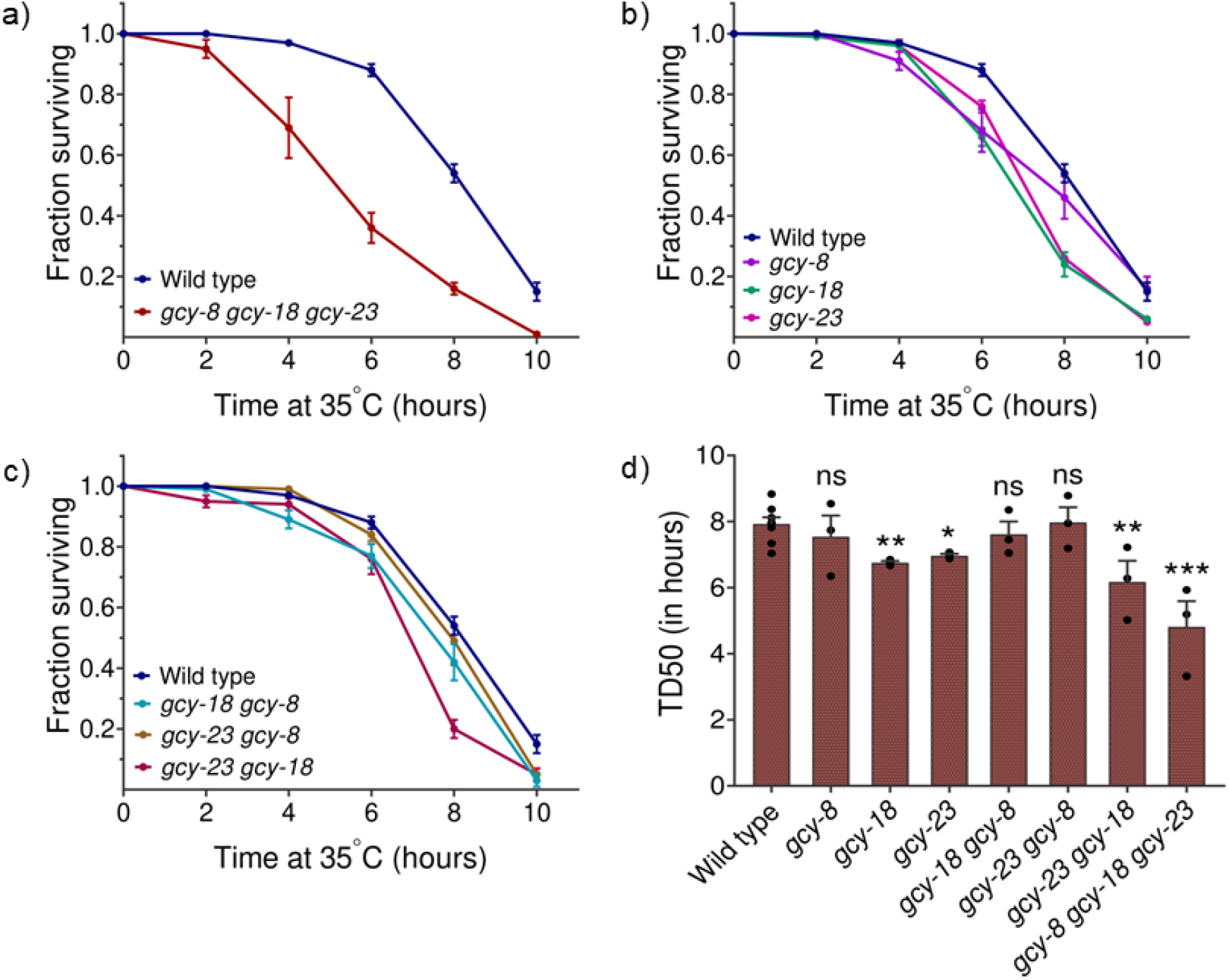
Effect of AFD-rGCs on the survival of adult worms under noxious temperatures. Survival fraction of rGC mutants compared to wild-type upon exposure to 35^°^C: **a)** Triple mutants (*gcy-8 gcy-18 gcy-23*), **b)** Single mutants (*gcy-8, gcy-18, gcy-23*) **c)** Double mutants (*gcy-18 gcy-8, gcy-23 gcy-8, gcy-23 gcy-18*). **d)** Mean *±* standard error of the half-survival time point (TD50) across genotypes. Data in all plots are means ± SEM from 3 biological replicates per group. Statistical comparisons were performed using an unpaired Student’s t-test against wild-type. ***p *<* 0.001, **p *<* 0.01, *p *<* 0.05; ns, not significant.

To quantify these differences in thermotolerance, we measured TD50 across all genotypes (shown in Fig. 1d). We observed that single mutants *gcy-18* and *gcy-23* show moderate but significant reduction in TD50 compared to wild-type worms, whereas, *gcy-8* mutants show no significant change. The effect is more pronounced in *gcy-23 gcy-18* double mutant, suggesting that some degree of functional overlap exists among these two rGCs. However, *gcy-8 gcy-18 gcy-23* triple mutant shows the most severe defect characterized by the lowest TD50 compared to all other groups, confirming that AFD-rGCs act together to support heat stress survival. The progressively lower TD50 in double and triple mutants highlights that while individual AFD-rGCs contribute to thermotolerance, their combined function is important. Together, these findings identify AFD-rGCs as key regulators of heat stress resistance, thus affecting the survival under extreme temperature conditions.

### Mutations in AFD-rGCs limit stress recovery in *C. elegans*

To determine whether AFD-rGCs influence both tolerance to and recovery from thermal stress, we examined their role in post-stress survival. We performed thermal stress recovery assays by exposing worms to 35^°^C for 4 h and subsequently monitored their lifespan (shown in Fig. 2). We first examined the *gcy-8 gcy-18 gcy-23* triple mutant, which showed a severely reduced lifespan after heat stress (Fig. 2a). This severe defect led us to investigate whether single and double mutants of rGCs also exhibit impaired post-stress survival. Further, on examining *gcy* single mutants (shown in Fig. 2b), we observed a significant reduction in survival than that of wild-type worms, thus indicating that the loss of any individual rGC can impair recovery. However, the overlap between the survival curves of *gcy-8, gcy-18* and *gcy-23* mutants suggests that the loss of function in each rGC impairs recovery to the same extent. This implies that no single rGC plays a more significant role in post-stress survival. We further observed that double mutants of AFD-rGCs do not show a significantly higher reduction in lifespan in comparison to their single mutant counterparts (Fig. 2c), which suggests that the loss of multiple rGCs do not have any additive effect on post-stress survival.

**Figure 2.**
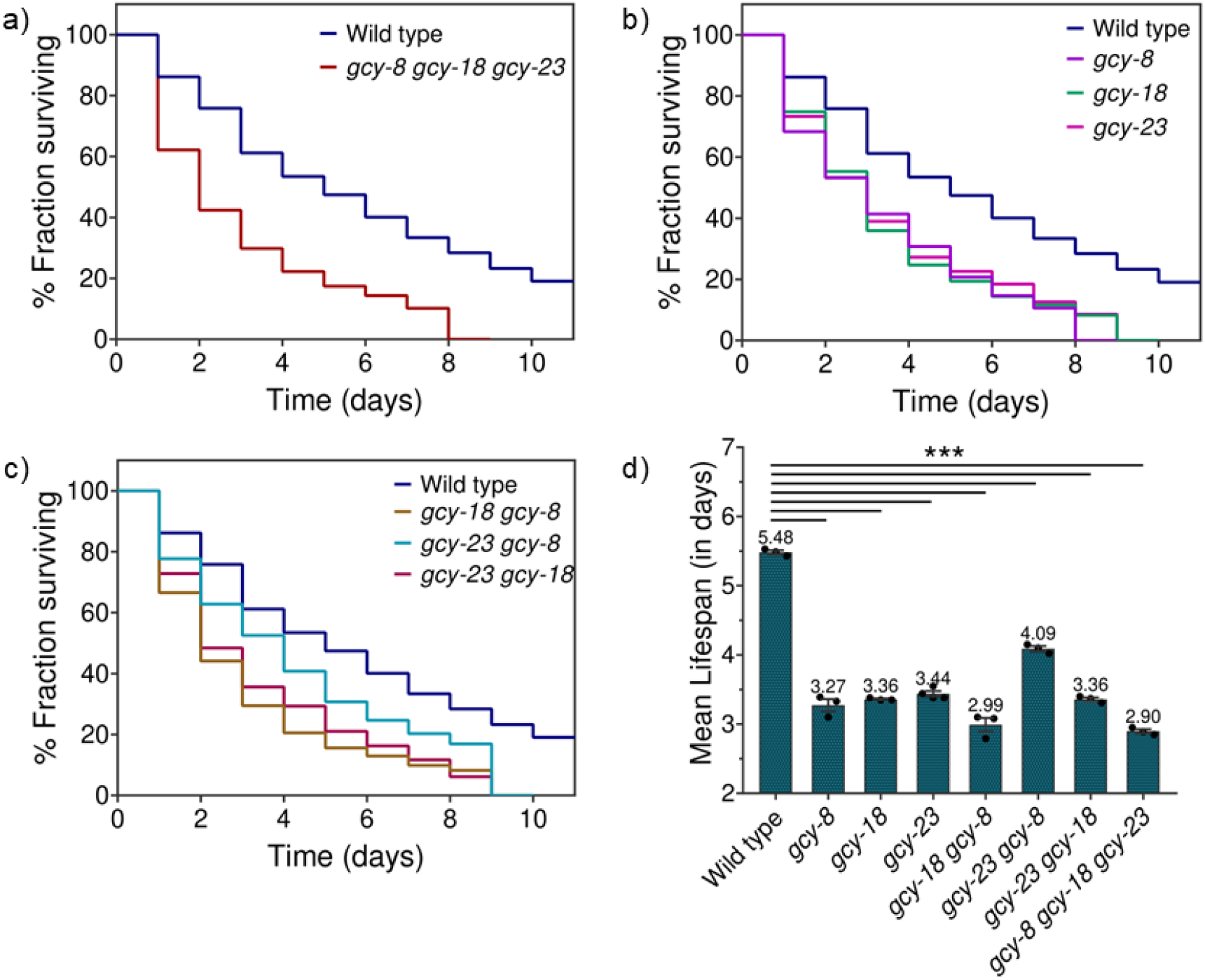
Effect of AFD-rGCs on post-stress survival in *C. elegans*. Percentage fraction survival following heat stress in rGC mutants compared to wild-type worms after exposure to 35^°^C for 4 h in : **(a)** Triple mutant (*gcy-8 gcy-18 gcy-23*), **(b)** Single mutants (*gcy-8, gcy-18, gcy-23*), **(c)** Double mutants (*gcy-18 gcy-8, gcy-23 gcy-8, gcy-23 gcy-18*). Adult hermaphrodites post heat shock were placed at 20^°^C and were monitored for survival. Data in (a), (b) and (c) are means from 3 biological replicates per group. **(d)** Mean lifespan comparisons across genotypes of *gcy-8, gcy-18, gcy-23* to wild-type animals post heat stress. Data are means ± SEM from 3 biological replicates per group. Statistical comparisons were performed using an unpaired Student’s t-test against wild-type. Error bars indicate the standard error of the mean (SEM) across biological replicates. ***p *<* 0.001, **p *<* 0.01, *p *<* 0.05; ns, not significant.

Similar to our thermotolerance results, where AFD-rGCs were essential for survival under prolonged heat exposure, we found that mutants lacking one or more rGCs have reduced lifespan post heat stress (Fig. 2a-c). Also, a comparison of mean lifespans across genotypes, shown in Fig. 2d further supports the trend, where wild-type worms survive the longest after heat stress (*∼* 5.48 days), however, all rGCs mutant groups show reduced mean lifespans with *gcy-8 gcy-18 gcy-23* triple mutant having the most severe defect. The shorter lifespan observed across different mutant genotypes of AFD-rGCs suggest that multiple rGCs work together to support stress recovery and their combined function is essential for long-term survival.

### Sensory AFD-rGCs facilitate a robust HSF-1 mediated stress response

The findings from physiological studies led us to investigate whether sensory perception via AFD-rGCs modulate the activation of protective mechanisms, such as the heat shock response (HSR), which is primarily regulated by the transcription factor HSF-1 and is known to promote chaperone induction under stress. In order to get deeper insights into this, we examined whether the induction of heat shock-induced chaperones was regulated by rGC signaling. Using quantitative RT–PCR, we analyzed the expression of HSF-1 target genes known to respond to heat shock in animals maintained at 20^°^C.

We first investigated the induction of *hsp-70* (encoded by the gene C12C8.1), a pivotal and well characterized HSR-induced, HSF-1 target gene in *C. elegans* [32]. To assess the role of AFD-rGCs in *hsp-70* regulation, wild-type and mutant adult animals grown at moderate population densities at 20^°^C with abundant bacterial food were transferred transiently for a heat shock (35^°^C for 1 h) and mRNA levels of *hsp-70* were measured one hour post-heat shock (Fig. 3a). We observed that the triple mutation in rGCs (*gcy-8 gcy-18 gcy-23*) caused a severe reduction in the heat shock–dependent accumulation of *hsp70* mRNA post heat shock. Among the single *gcy* mutants in Fig. 3a, the significantly reduced expression of *hsp-70* in *gcy-18* mutant suggests that *gcy-18* plays a major role in HSR regulation, while *gcy-8* and *gcy-23* may have redundant or supporting functions. The further reduction observed in the *hsp-70* accumulation in *gcy-18 gcy-23* double mutant reinforces the idea that *gcy-18* is crucial for HSR induction through *hsp-70*, with *gcy-23* playing a contributing role.

**Figure 3.**
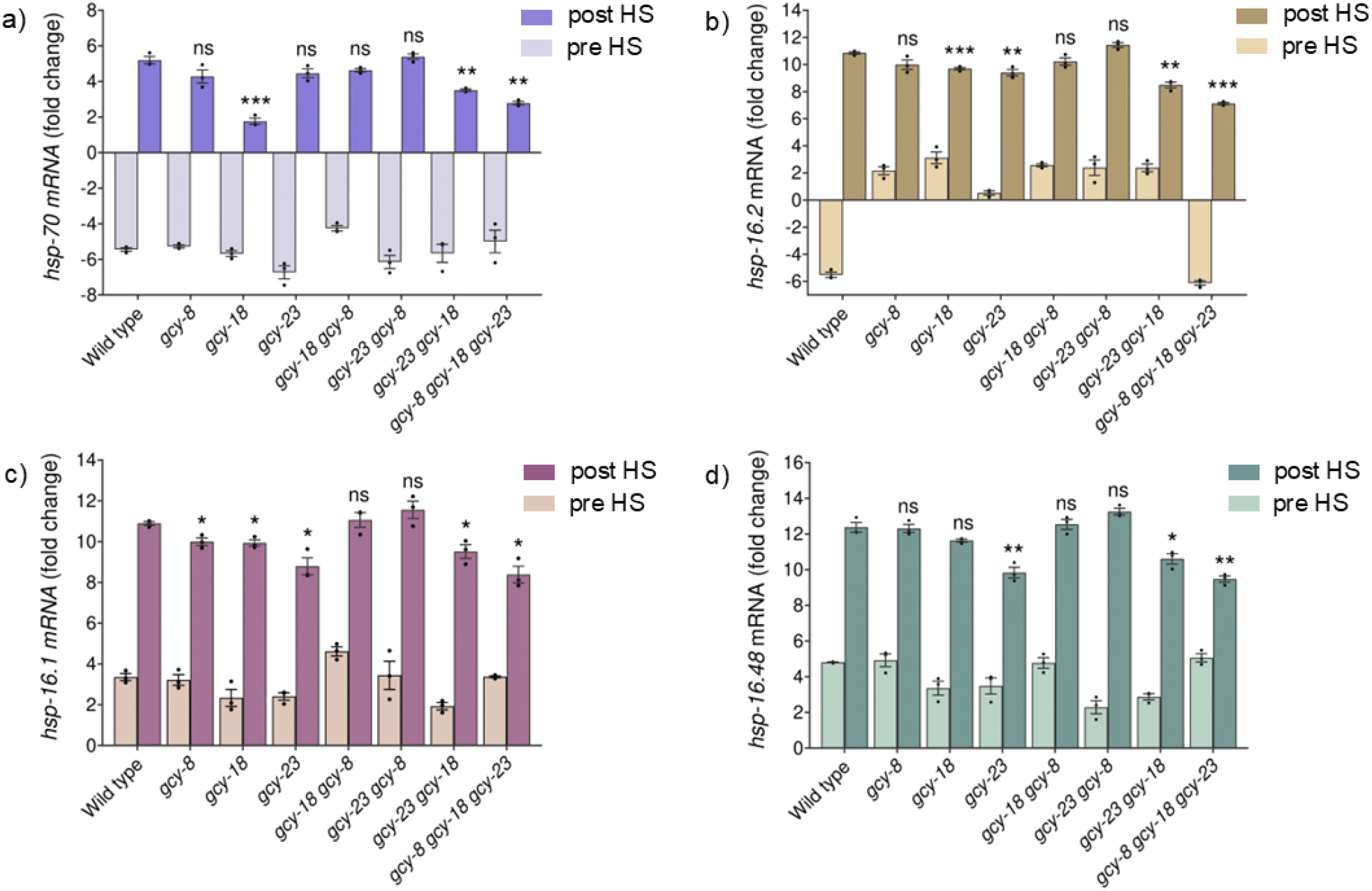
Transcript level changes of heat shock proteins involved in the HSR pathway before and after heat shock at 35^°^C. Fold change in mRNA levels of **(a)** *hsp-70*, **(b)** *hsp-16*.*2*, **(c)** *hsp-16*.*1*, **(d)** *hsp-16*.*48* in wild-type, single, double and triple *gcy* mutants. Expression levels were measured in worms collected 1 h post heat shock and comparisons are performed to post-heat shock levels in wild-type. Data represent mean ± SEM from three independent biological replicates. Statistical comparisons were performed using Welch’s t-test. Asterisks indicate significance relative to wild-type worms post heat shock (***p *<* 0.001, **p *<* 0.01, *p *<* 0.05; ns, not significant).

Next, to further confirm that rGC mutations suppress cytosolic proteostasis by downregulating HSF-1 regulated genes, we examined the mRNA expression of several small HSPs. Small heat shock proteins (sHSPs) function as ATP-independent chaperones that prevent protein aggregation by transiently sequestering misfolded proteins until they can be refolded by the ATP-dependent *hsp-70* machinery [33, 34]. To uncover the effect of *gcy* mutations on the induction of HSR, we focused on the *hsp-16* family, as these chaperones are typically undetectable under normal physiological conditions, neither with Northern blot nor immunostaining [35, 36]. However, these sHSPs play a crucial role in the mediation of heat resistance upon heat stress [37–41], that makes them ideal candidates for assessing disruptions in HSR signaling and proteostasis mechanisms.

Given the crucial role of sHSPs in maintaining proteostasis under stress, we compared the mRNA levels of *hsp-16*.*2, hsp-16*.*48* and *hsp-16*.*1* before and after heat shock. In wildtype animals, all three sHSPs were upregulated post-heat shock, however, mutations in *gcy* genes significantly impair this response (Fig. 3b-d). First, we observed that mRNA levels for *hsp-16*.*2, hsp-16*.*48* and *hsp-16*.*1* were significantly suppressed in *gcy-8 gcy-18 gcy-23* triple mutant, strongly suggesting that cytoplasmic stress responses rely on receptor-based signaling in AFD neurons under noxious thermal stress. On examining single mutants of rGCs, we found that *gcy-23* mutant has reduced mRNA levels across all three sHSPs, indicating an important role for *gcy-23* in regulating their transcription. Similarly, *gcy-18* mutants showed suppressed induction of *hsp-16*.*2* and *hsp-16*.*1* genes, that suggests its essential function in HSR activation. As shown in Fig. 3, further analysis of rGC double mutants revealed that only the *gcy-23 gcy-18* double mutant exhibits a significant reduction in sHSPs expression compared to wild-type animals post-heat shock. This highlights the important role of both, *gcy-23* and *gcy-18*, in activating the HSR and suggests a cooperative regulation of sHSPs by these AFD-rGCs. As observed, the expression of sHSPs in the double mutant *gcy-23 gcy-18* was, in some cases, reduced to a greater extent than in either single mutant, suggesting that these genes likely function in parallel or within the same pathway to modulate HSR activation. However, *gcy-8* seems to be an exception. Not only did loss of function in *gcy-8* alone lead to no alteration in the post heat shock induction of *hsp*s in most cases, double mutants of *gcy-8* with either *gcy-18* or *gcy-23* was observed to suppress the phenotypic reductions found in the corresponding single mutants. This observation points to complex inter-dependencies within the rGC signaling, where feedback mechanisms potentially at the level of transcriptional regulation adjust the output to maintain signaling homeostasis. Further, since pre-heat shock mRNA levels remained low across all genotypes, these effects specifically originate from impaired heat-induced transcription rather than defects in basal expression. Overall, these results suggest that receptor-based signaling in AFD neurons is crucial for cytoplasmic stress responses, with *gcy-23* and *gcy-18* serving as primary regulators of sHSP induction to maintain proteostasis under thermal stress.

### HSR activation defects in rGC mutants are not merely due to a delayed onset

To determine whether the reduced expression of heat shock induced genes in AFD-rGC mutants result from a delayed response, we analyzed the temporal dynamics of *hsp-70, hsp-16*.*2, hsp-16*.*1* and *hsp-16*.*48* mRNA levels over a 6 h period following 1 hour of heat shock at 35^°^C. In wild-type animals, we observed a rapid induction for all *hsp*s, reaching their peak expression 1 h post-heat shock, *i*.*e*. at the 2 h timepoint, before gradually declining (Fig. 4). However, the *gcy-8 gcy-18 gcy-23* triple mutant (shown in Fig. 4a-d) showed significantly lower or not significantly different *hsp* induction across all the time points, therefore indicating a persistent decline in the HSR induction rather than a mere delay. By comparison, maximal levels in *gcy-8 gcy-18 gcy-23* triple mutants were lower by *∼*40-50% compared to wild-type animals at 1 h post-heat shock (2 h time point), thus reflecting a diminished transcriptional response in rGCs mutant. We found that beyond the peak expression, *hsp* mRNA levels in the *gcy* triple mutant exhibited altered decay kinetics, with a more rapid post-peak decline observed for specific genes such as *hsp-16*.*1* and *hsp-16*.*48*, compared to wild type following heat shock (Fig. 4c–d). Together, these findings suggest that AFD-rGCs not only modulate the initial magnitude of the heat shock response but also influence its duration, likely by regulating HSF-1 mediated transcription. The impaired induction and, for certain *hsp* genes, a more rapid post-peak decay of *hsp* levels in *gcy* mutants suggest a broader role for AFD-rGC signaling in maintaining proteostasis under thermal stress.

**Figure 4.**
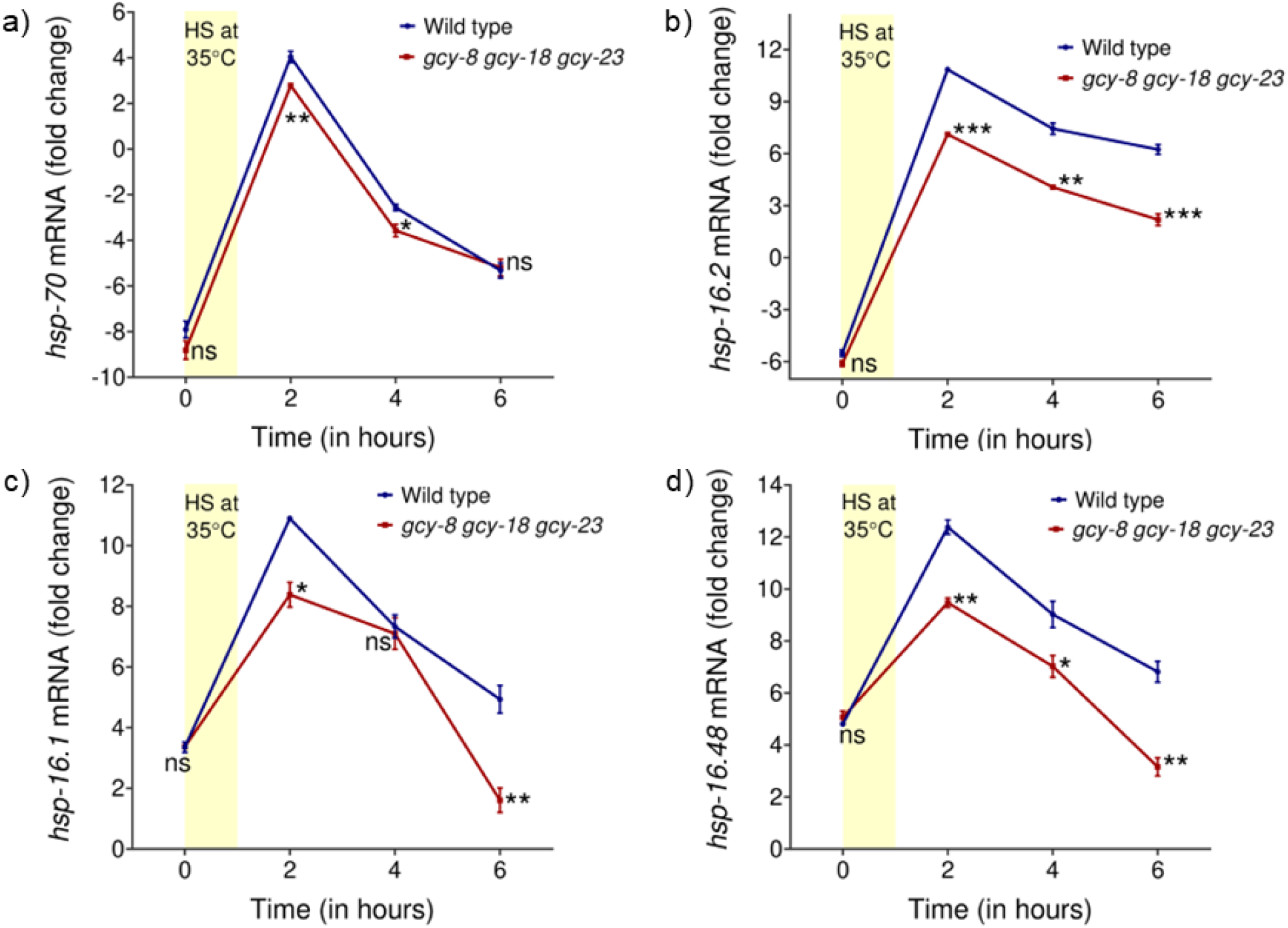
AFD-rGCs regulate heat shock protein expression dynamics following thermal stress. Transcript levels of **a)** *hsp-70*, **b)** *hsp-16*.*2*, **c)** *hsp-16*.*1*, **d)** *hsp-16*.*48* in wild-type and *gcy-8 gcy-18 gcy-23* triple mutant at pre and post-heat shock time points (0, 2, 4 and 6 h). The shaded yellow region represents the heat shock period (35^°^C for 1 h). All values represent mean *±* SEM from at least three biological replicates. Statistical significance was determined using Welch’s t-test comparing levels in wild-type to *gcy* triple mutant at each time point. ***p *<* 0.001, **p *<* 0.01, *p *<* 0.05, ns, not significant.

### Impaired physiological responses in rGC mutants result from downregulation of small heat shock chaperones

Among the mRNA levels of *hsp-16* genes analyzed (Fig. 3b-d), *hsp-16*.*2* appears to be most significantly affected by *gcy* mutations, with the strongest downregulation observed in their triple mutant (shown in Fig. 3b). When examining thermotolerance trends across wild-type and *gcy* mutant genotypes (Fig. 1d), we observed that the ability of animals to withstand heat stress, as indicated by TD50 values, closely follows the trend of *hsp-16*.*2* mRNA expression across different *gcy* mutants. *hsp-16*.*2* is a well-characterized sHSP important for maintaining protein homeostasis under thermal stress [26, 29, 36, 40] and has been extensively studied as a stress response marker [42–44]. Therefore, its downregulation in rGC mutants suggest a potential impairment in cellular stress response mechanisms. Prior studies have shown that adult *C. elegans* with reduced *hsp-16*.*2* expression exhibit shorter lifespans and diminished survival under thermal stress [1, 3, 39]. This aligns with our observations, where the loss of AFD-rGC genes appear to compromise thermotolerance, possibly by reducing the induction of sHSP, *hsp-16*.*2*, that is normally activated by heat stress to regulate stress-responsive gene expression. While these findings highlight a strong association between the reduced thermotolerance or stress resistance in rGC mutants to decreased accumulation of *hsp-16*.*2* (Fig. 1d and Fig. 3b), it is likely that additional factors contribute to the observed phenotype. Other sHSPs along with broader cellular stress response pathways may play a synergistic role in contributing to the observed thermotolerance behavior in *gcy* mutants.

Furthermore, as previously shown in Fig. 2d, our findings indicate that mutations in AFD-rGCs result in significant reduction in lifespan of animals following heat shock treatment, which coincides with the altered trend of *hsp-12*.*6* mRNA expression post-heat shock (Fig. S1a-b). While *hsp-12*.*6* lacks chaperone activity in vitro, it has been reported to act downstream of DAF-16 [45, 46], the primary transcription factor regulating lifespan extension under reduced insulin/IGF-1 signaling [6, 46, 47]. More recent studies further highlight *hsp-12*.*6* as a positive effector of both lifespan and reproduction in *C. elegans* [48]. The observed downregulation of *hsp-12*.*6* post heat shock in *gcy* mutants compared to wild type (Fig. S1), along with their shortened mean lifespan following heat stress, suggests that AFD-rGC signaling may affect the other intersecting arm of HSR mechanism [47], downstream of DAF-16 to regulate stress adaptation.

### Thermosensory rGC signaling modulates inter-tissue communication

We next examined whether the decreased accumulation of HSPs in rGC mutants reflects a selective reduction in neuronal tissue or results from impaired induction of these HSPs in distant tissues upon exposure to high temperature, suggesting a role for inter-tissue communication. While our HSPs transcript level study suggest that *hsp-70* and other sHSPs are affected in *gcy* mutants, *hsp-16*.*2* emerged as the one most significantly impacted among them. The mRNA levels of *hsp-16*.*2* were markedly downregulated (Fig. 3b) and its expression pattern closely mirrored the observed thermotolerance defects (Fig. 1d) in rGC mutants. Given this strong correlation and the established role of *hsp-16*.*2* as a key stress-responsive marker [39, 43, 44, 49], we focused on this particular sHSP to explore whether AFD-rGCs contribute to inter-tissue communication during heat stress. To investigate this, we used worms expressing GFP under the regulation of the *hsp-16*.*2* promoter (strain TJ3001), which is a well-characterized HSF-1 target predominantly expressed in the intestine [1, 42]. This reporter strain carries the *hsp-16*.*2* promoter sequence fused to the GFP gene from jellyfish [50], allowing real-time visualization of *hsp-16*.*2* activation. Although this reporter does not encode the HSP-16.2 protein product itself, previous studies have demonstrated that GFP expression from this reporter parallels endogenous *hsp-16*.*2* gene upregulation, indicating that *Phsp-16*.*2::GFP* is a reliable reporter for *hsp-16*.*2* induction. Our mRNA analysis of *gfp* expression in this reporter strain further validated its use as a reliable proxy for monitoring changes in HSP-16.2 induction, as its expression pattern closely followed the time-dependent trend observed for endogenous *hsp-16*.*2* levels (Fig. S2).

To further investigate the role of AFD expressed rGCs in regulating HSP-16.2 expression, we quantified fluorescence intensity from the *Phsp-16*.*2::GFP* reporter in wild-type (Fig. 5a) and *gcy-8 gcy-18 gcy-23* triple mutants, with and without heat shock (35^°^C, 1 h). We observed the baseline fluorescence levels remained low in both genotypes under non-stress conditions. However, post heat shock, the GFP levels in *gcy* triple mutants did not rise to the same extent as in wild-type worms (Fig. 5b), indicating that AFD-rGCs play an important role in amplifying HSP-16.2 expression during the heat shock response. To dissect the individual contribution of each rGC, we analyzed fluorescence intensity in single and double *gcy* mutants of *gcy-8, gcy-18* and *gcy-23* before (Fig. S4) and after heat shock (Fig. 5c). We observed that loss of individual rGCs (single *gcy* mutants) led to a significant reduction in HSP-16.2 expression compared to wild type after heat stress (Fig. 5c), therefore indicating that each rGC contributes to the heat shock–evoked activity of AFD neurons. However, as shown in Fig. 5c, fluorescence intensity levels across all rGC double mutants remained lower than wild type but were often higher than the more severely affected single mutants *i*.*e. gcy-8, gcy-18* and *gcy-23*, suggesting partial functional compensation when multiple rGCs are simultaneously disrupted. These findings suggest that AFD-expressed rGCs collectively modulate post-stress translational responses, highlighting their integrated role in shaping downstream responses after stress exposure.

**Figure 5.**
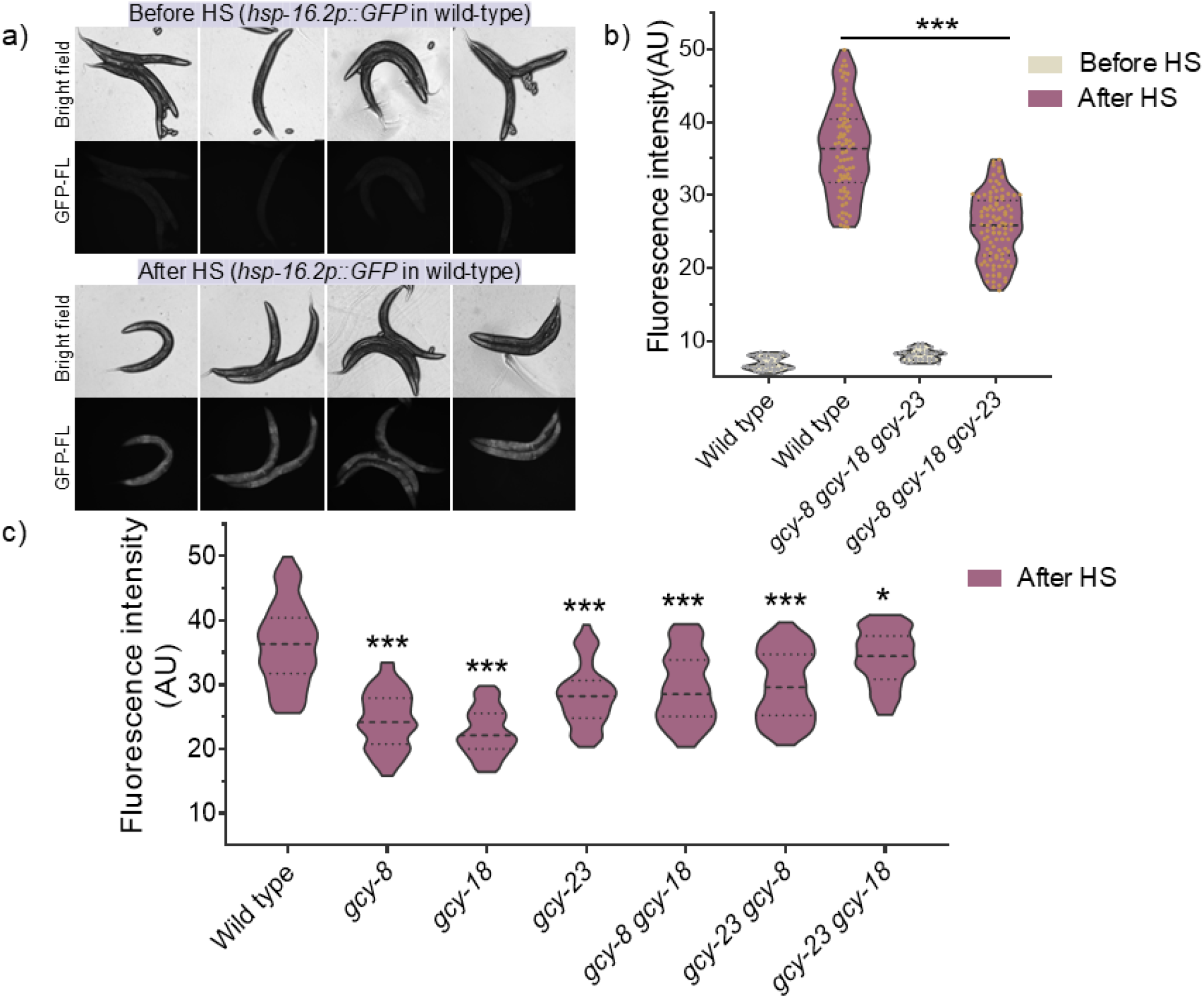
AFD-rGCs modulate HSP-16.2 expression in non-neuronal tissues after heat stress. **a)** Representative Bright field and GFP fluorescence (GFP-FL) images of *hsp-16*.*2p::GFP* reporter in wild-type animals before and 5 h after heat shock (35^°^C for 1 h). Imaging was performed at exposure times of 70 ms for brightfield and 300 ms for GFP-FL. **b)** Quantification of GFP fluorescence intensity in wild-type and *gcy-8 gcy-18 gcy-23* triple mutant animals before and after heat shock. Each dot in the plot represents the fluorescence intensity of an individual worm. **c)** Fluorescence intensity quantification after heat shock in single and double mutants of AFD-rGCs, *gcy-8, gcy-18* and *gcy-23*. Each experiment was repeated at least three times, with 30–40 worms imaged per condition. Fluorescence intensity is shown in arbitrary units (AU). Statistical significance was determined using Student’s t-test comparing levels in wild-type after heat stress to *gcy* mutants; *p *<* 0.05, ***p *<* 0.001.

Overall, these findings support the idea that AFD neurons do not merely function as thermosensors but also mediate inter-tissue signaling to regulate stress-responsive gene expression. The dampened HSP-16.2 induction in *gcy* mutants indicates that rGC activity within AFD neurons is essential for modulating the magnitude of the heat shock response in peripheral tissues, such as the intestine. This further reinforces the hypothesis that the heat shock response in *C. elegans* is not entirely cell-autonomous under noxious conditions but relies on neuronal input for proper activation.

## SUMMARY AND DISCUSSION

The thermosensory AFD neurons in *C. elegans* are integral to the organism’s ability to detect and respond to temperature changes. These neurons enable the worm to sense minute temperature fluctuations, facilitating behaviors such as thermotaxis and avoidance of noxious heat [2, 51, 52]. Within these neurons, three *gcy-8* -, *gcy-18* - and *gcy-23* -encoded rGCs are specifically expressed and localized to the specialized sensory endings [15, 53].

Mutants lacking all these three rGCs exhibit abolished temperature-evoked currents in AFD, therefore demonstrating that these rGCs are necessary for thermotransduction and serve as primary thermosensors in these neurons [19]. While previous studies have examined the contributions of AFD-expressed rGCs to thermotransduction under moderate temperature conditions (15–25^°^C) [15, 16, 54, 55], our understanding of their specific functions in sensing and responding to noxious thermal stress remains limited.

To address this gap and recognizing that extreme temperatures pose a severe challenge to organismal survival, we systematically investigated the contributions of GCY-8, GCY-18, and GCY-23 to AFD-mediated responses under noxious temperature conditions (*>*30^°^C). Through a combination of genetic and physiological analyses across single, double and triple mutants, we uncover both distinct and overlapping roles of these rGCs in thermotransduction. Our findings establish that AFD-rGCs play an important role in survival under heat stress, with each contributing distinctly to the activation of cellular stress response mechanisms in *C. elegans*.

Since rGCs play a key role in thermotransduction, we hypothesized that reduced signaling via these rGCs might impair the physiological ability of worms to withstand elevated temperatures. Consistent with this, we found that a triple mutation in rGCs (*gcy-8 gcy-18 gcy-23*) compromises the ability of worms to tolerate thermal stress (Fig. 1). Our analysis further indicated a functional overlap between *gcy-18* and *gcy-23*, underscoring their cooperative role in heat resistance. Additionally, we found AFD-rGCs to be essential not only for survival under thermal stress but also for post-stress recovery, as observed by the severely reduced lifespan of triple mutants and defects observed in single and double mutants after heat exposure (Fig. 2). Given these physiological impairments, we reasoned that the underlying cause might lie in the reduced activation of cellular stress response pathways, which are crucial for mitigating damage under extreme conditions. Since the HSR is a primary protective mechanism against thermal stress, we next examined whether mutations in AFD-rGCs disrupt the transcriptional regulation of HSPs, potentially explaining the observed defects.

Our analysis suggests that AFD neuron expressed rGCs play important contribution in coordinating heat shock responses at the molecular level. Following heat shock (35^°^C, 1h), we observed a significant reduction in *hsp-70* mRNA levels across *gcy* mutants, with the most severe impairment in the triple mutant. Further, genotype comparison for *hsp-70* levels across single and double mutants of *gcy* highlighted *gcy-18* as a key regulator of HSR induction, while *gcy-23* played a supporting role. Intriguingly, ATP-independent holdase chaperones, particularly small HSPs from the *hsp-16* family known for their rapid induction under thermal stress [37–41] also showed impaired activation in these mutants. This disruption in the HSF-1 mediated stress response suggests that AFD-rGCs are not merely sensors of temperature but also critical modulators of cellular defense pathways. Our findings indicate that receptor-based signaling in AFD neurons is essential for cytoplasmic stress responses, with *gcy-18* and *gcy-23* serving as primary regulators of sHSP induction to maintain proteostasis under thermal stress. Furthermore, time-dependent analyses of mRNA levels revealed that these rGCs influence both the magnitude and duration of the HSR, likely through HSF-1-mediated transcription. Importantly, the defects observed in *gcy* mutants were not merely due to a delayed response but reflected an inability to mount robust and lasting defense against thermal stress. Extending this connection, we also observed altered expression of *hsp-12*.*6*, a small HSP that lacks chaperone activity but functions downstream of DAF-16. Given its established role in longevity and reproduction [46, 47], the reduced *hsp-12*.*6* expression in *gcy* mutants, coupled with their shorter post-heat shock lifespan suggests that AFD-rGC signaling may intersect with the DAF-16 pathway. This highlights a broader link between thermosensory signaling and lifespan regulation, indicating that AFD neurons may coordinate stress adaptation through multiple molecular pathways.

The marked downregulation of *hsp-16*.*2* mRNA in *gcy* mutants, along with its expression pattern mirroring thermotolerance defects, highlights its importance as a stress response marker to study regulation of HSR by AFD-rGCs. Beyond the role of AFD-rGCs in thermosensation, our findings reveal that AFD-rGCs also differentially contribute to intertissue communication during heat stress. Using the *Phsp-16*.*2::GFP* reporter strain, we found that the reduced accumulation of HSP-16.2 in rGC mutants was not only limited to neuronal tissue but also led to impaired induction in distant tissues upon heat exposure. These findings reinforce the idea that the HSR in *C. elegans* is not entirely cell-autonomous but instead relies on neuronal input for proper activation. Therefore, AFD neurons function beyond thermosensation and actively mediate inter-tissue communication to coordinate stress-responsive gene expression (Fig. 6).

**Figure 6.**
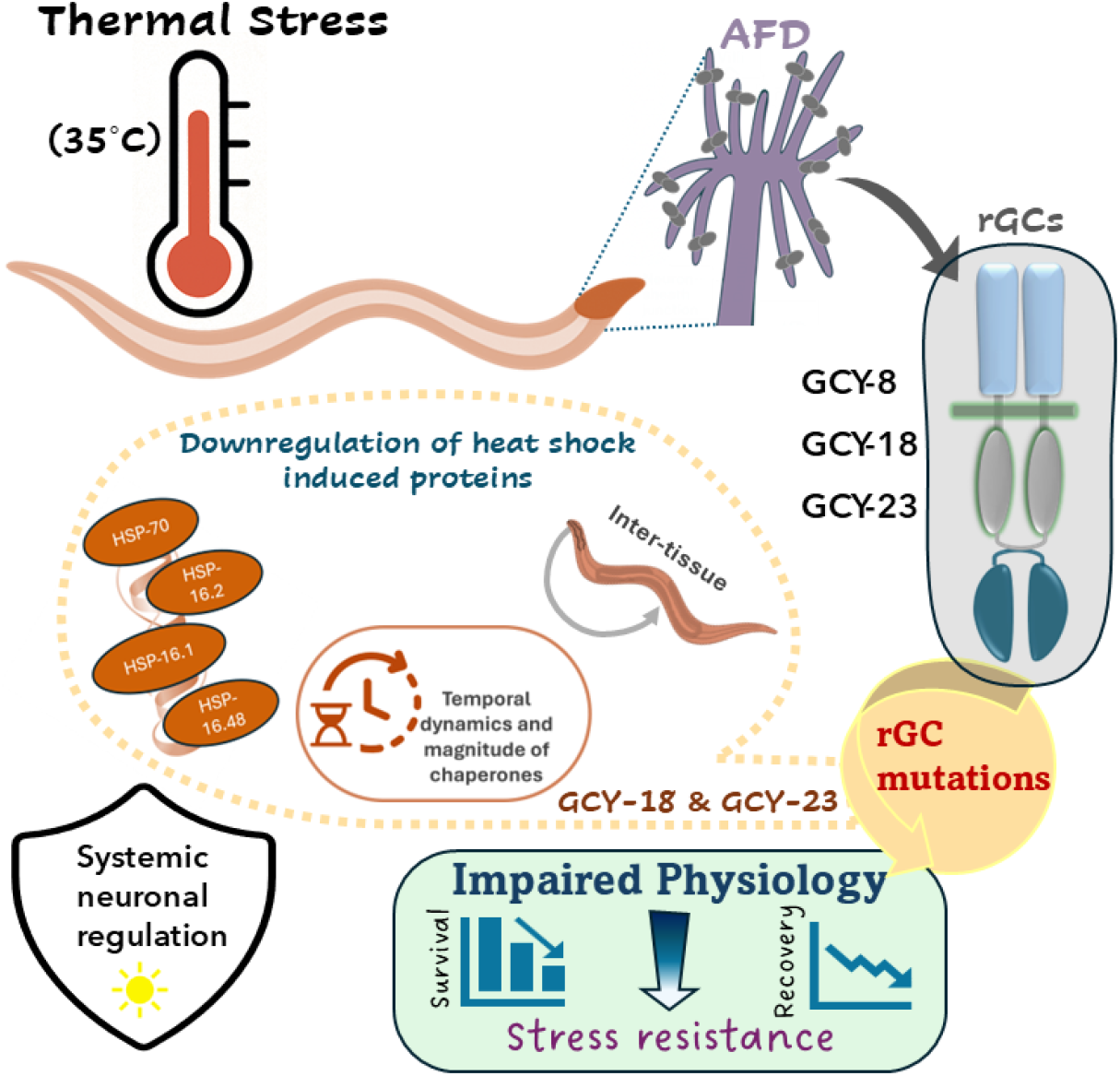
AFD-rGCs mediate systemic regulation of the heat shock response in *C. elegans*. Exposure to thermal stress (35^°^C) activates AFD thermosensory neurons, which signal through rGCs— GCY-8, GCY-18 and GCY-23 to modulate organism-wide heat shock responses. Mutations in GCY-18 and GCY-23 alter the temporal dynamics and magnitude of chaperone induction, including small heat shock proteins (HSP-16.1, HSP-16.2, HSP-16.48) and HSP-70 therefore leading to impaired proteostasis. This systemic modulation impacts stress resistance, survival and post-stress recovery, thus highlighting a neuro-sensory control that governs the cellular stress responses.

More broadly, our results indicate that different rGCs contribute distinctly to thermosensory responses under noxious temperature conditions, with *gcy-18* and *gcy-23* primarily governing transcriptional induction of HSR. Given the presence of rGCs as thermosensitive proteins in both *C. elegans* and mammals, their functional diversity suggests an evolutionarily conserved role in thermosensory signaling across phyla. These findings open exciting possibilities for exploring how thermosensory networks regulate cellular stress responses in other organisms, potentially unveiling fundamental principles of sensory-driven stress adaptation.

## MATERIAL AND METHODS

### Nematode strains

All *C. elegans* strains were cultivated on standard nematode growth medium (NGM) plates seeded with *E. Coli* OP50 strain and maintained at 20^°^C as described in [56]. The strains below were obtained from the Caenorhabditis Genetics Center (CGC): wild type N2 Bristol, IK800[*gcy-8(oy44)*], IK429[*gcy-18(nj38)*], IK427[*gcy-23(nj37)*], IK597[*gcy-23(nj37) gcy-8(oy44) gcy-18(nj38)*], TJ3001[*hsp-16*.*2p::GFP::unc-54 + Cbr-unc-119(+)*], EG7935[*unc-119(ed3); eft-3p::tdTomato::H2B::unc-54 3’UTR + Cbr-unc-119(+)*]. The double mutant *gcy* strains were provided by Prof. Ikue Mori (Nagoya University, Japan). A complete list of strains used in this study are provided in Table S1.

### Thermotolerance assays

To obtain a synchronized population, 3-4 gravid hermaphrodites were transferred onto seeded NGM plates at room temperature and allowed to lay 20–30 eggs per plate. After approximately 3 hours, the gravid worms were removed to maintain synchronous development of the progeny, which were kept at 20^°^C for *∼* 60 h. On day 3, plates with young adult worms were transferred to a pre-equilibrated incubator at 35^°^C for heat shock treatment. Stacking of plates was avoided during heat shock. For technical replication, minimum three plates were removed from the incubator at 2 h intervals after heat stress durations ranging from 2 to 10 h. The plates were left at room temperature for a short recovery period of 15 min. Worms were examined on each plate using a platinum wire and those that did not respond to touch with the platinum pick were scored as dead. All assays were performed with total 60–100 animals at least three times.

### Stress recovery assays

For thermorecovery assays, synchronized day 3 adult worms (20–30 per plate) on *E. Coli* OP50 seeded plates were subjected to heat shock at 35^°^C for 4 h in an incubator. Survival was then scored at 24 h intervals following heat shock until the entire population was dead. To prevent interference from progeny, animals were transferred to new plates every day for the first 4 days of adulthood and every 2-3 days thereafter. Worms were scored as dead in the absence of pharyngeal pumping and response to touch with a platinum pick. Dead worms with vulval rupture, worms that died from climbing up the side of the plates and missing worms were censored from the analysis.

### Heat shock treatment

For mRNA expression studies, bulk worm samples were heat shocked on 60 mm *E. Coli* OP50 seeded NGM plates by transferring them to a pre-equilibrated incubator set at 35^°^C. Worms were exposed to heat shock for a duration of 1 h, after which they were collected in M9 buffer following the completion of the specified recovery periods.

### RNA extraction and cDNA synthesis

RNA was extracted from approximately 600 adult worms, which were washed 2–3 times to remove bacteria before lysis and homogenization in 1 mL of Trizol extraction reagent. The samples were vortexed for 30 s, snap-frozen in liquid nitrogen, then thawed at 37°C.

This cycle was repeated three times. Tubes with worm sample were then stored at −80^°^C until further processing. Before RNA extraction, samples were thawed and left at room temperature for 5 min. Worms were then vortexed for 2 min, with 2 min intervals on ice, for a total of four cycles. Following this, 300 *µ*L chloroform was added to Trizol mixture, shaken for 15 sec by hand, and allowed to stand at room temperature for 4 min. Samples were then centrifuged at 13,000 rpm for 15 min at 4^°^C. The aqueous phase was then transferred to fresh tubes and gently mixed with 500 *µ*L of isopropanol by inverting the tubes, followed by incubation at room temperature for 15 min. The sample in the tubes was then centrifuged at 12,000 rpm for 15 min at 4^°^C, resulting in the formation of a small white RNA pellet at the bottom of the tube. After removing the supernatant, the pellet was washed twice with ice cold 75% EtOH, followed by centrifugation at 4000 rpm at 4^°^C for 5 min. Once nearly all the ethanol had evaporated, the dried pellet was resuspended and dissolved in 40 µL of 1× TE buffer.

Next, RNA was purified using DNase I digestion (ThermoFisher RapidOut DNA Removal Kit) to remove genomic DNA and steps were followed as per manufacturer’s instructions. Finally, 1 mg of total RNA was reverse-transcribed into cDNA using the BioRad iScript cDNA synthesis kit, as per the manufacturer’s instructions.

### qRT-PCR and gene expression analysis

Relative mRNA expression was assessed via real-time quantitative PCR using iTaq Universal SYBR Green Supermix (BioRad) on a QuantStudio 5 Real-Time PCR System (Thermo Fisher Scientific). Reactions were performed in a total volume of 10 *µ*L using a 96-well plate. The fold change in mRNA levels between pre- and post-heat shock samples was calculated using the 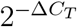 method. Gene expression was normalized to the house-keeping gene *cdc-42*, which was selected as an internal control due to its stable expression across stress conditions compared to other commonly used reference genes, such as *act-1* and *ire-1. C*_*T*_ values were obtained in triplicate for each sample (technical replicates), and each experiment was repeated at least three times (biological replicates) to determine Δ*C*_*T*_. Reverse transcriptase minus controls were included to confirm the absence of genomic DNA contamination and most set of primers were designed to target exon–exon junctions to further minimize amplification of any residual genomic DNA. All primer sequences used for qRT-PCR are listed in Table S2.

### Genotype assessment

The genotype of the crossed strain of single, double and triple mutant of *gcy-8(oy44) IV, gcy-18(nj38) IV* and *gcy-23(nj37) IV* with TJ3001 was determined by PCR using primer pairs designed to detect both wild-type and mutant alleles, as listed in Table S2 (Fig. S3). Individual adult worms were lysed for genotyping in 1× lysis buffer (50 mM KCl, 10 mM Tris pH 8.2, 2.5 mM MgCl_2_, 0.45% NP-40, 0.45% Tween 20, 0.5% Gelatin (2%)). To prepare the lysis mixture, 2.5 *µ*L of 20 mg/mL Proteinase K was added to 47.5 *µ*L of lysis buffer. Individual worms were lysed in 5 *µ*L of this mixture, followed by incubation at 65^°^C for 1 h and heat inactivation at 99^°^C for 7 minutes. PCR amplification was performed using 3 *µ*L of the lysate under the following cycling conditions: initial denaturation at 95^°^C for 2 min, followed by 35 cycles of denaturation at 95^°^C for 30 sec, annealing at T_*m*_ (*∼*60^°^C) for 30 sec and extension at 72^°^C for 2 min, with a final extension at 72^°^C for 10 min.

### Fluorescence imaging and quantification of fluorescence intensity

To assess the induction of stress reporter, transgenic worms were given heat shock by transferring them from standard growth conditions to a pre-equilibrated incubator set at 35^°^C for 1 h. Following heat shock, worms were allowed to recover at room temperature for approximately 5 h before imaging to capture peak induction levels of the reporter. For imaging, worms were immobilized on 2% agarose pads with 10 *µ*L of 5 mM levamisole and imaged immediately. Brightfield and fluorescence images were acquired using a Leica DMI8 inverted microscope equipped with a 10× objective, a DFC7000T camera and LAS X software. The emission wavelengths for the brightfield and fluorescence channels were 550 nm and 527 nm, respectively, with exposure times specified in the figure legends.

Images were acquired in binning mode (1×1; 1920×1440 resolution) and analyzed using FIJI [57]. Fluorescence intensity was quantified by manually outlining regions of interest (ROIs) corresponding to individual worms. For each worm, the mean GFP fluorescence intensity was measured and normalized to its area to account for size variations. Background fluorescence was determined by averaging the intensity of multiple background ROIs and subtracted from all measurements to obtain the corrected fluorescence intensity per worm. For the ease of comparison across samples, the normalized fluorescence intensity values were multiplied by an arbitrary factor of 10^5^.

### Quantification and statistical analysis

For thermotolerance analysis, comparisons of means and standard errors were supplemented with p-values obtained using two-tailed Student’s t-test. Mean lifespan for stress resistance assays was calculated using OASIS 2 [58], an online survival analysis tool that applies Kaplan-Meier estimation and the Log-rank (Mantel-Cox) test. For all other comparisons, statistical tests were selected as appropriate and are specified in the figure legends. One-way ANOVA with Tukey’s post hoc test was used for pairwise group comparisons, while a two-tailed Student’s t-test was applied where indicated. Fluorescence quantification was performed with a total of 100–120 worms across treated and untreated groups, with minimum three independent trials. All statistical analyses were performed using GraphPad Prism or Microsoft Excel, as specified in the figure legends.

## DATA AVAILABILITY

All study data are included in the article and/or supporting information.

## ACKNOWLEDGMENTS

R.S. acknowledges support for this work from ANRF (formerly known as Science and Engineering Research Board or SERB) through the Startup Research Grant (No. SRG/2019/000726) and the SERB Power Grant (No. SPG/2021/002732).

## DECLARATION OF INTERESTS

The authors declare no competing interests.

## Notes

### Competing Interest Statement

The authors have declared no competing interest.

