## Supplementary Information for "Guanylyl cyclase signaling in AFD neurons regulates systemic stress resilience in *Caenorhabditis elegans*"

**This PDF file includes:**

Figures S1-S4

Table S1-S3

SI References

### SUPPORTING INFORMATION

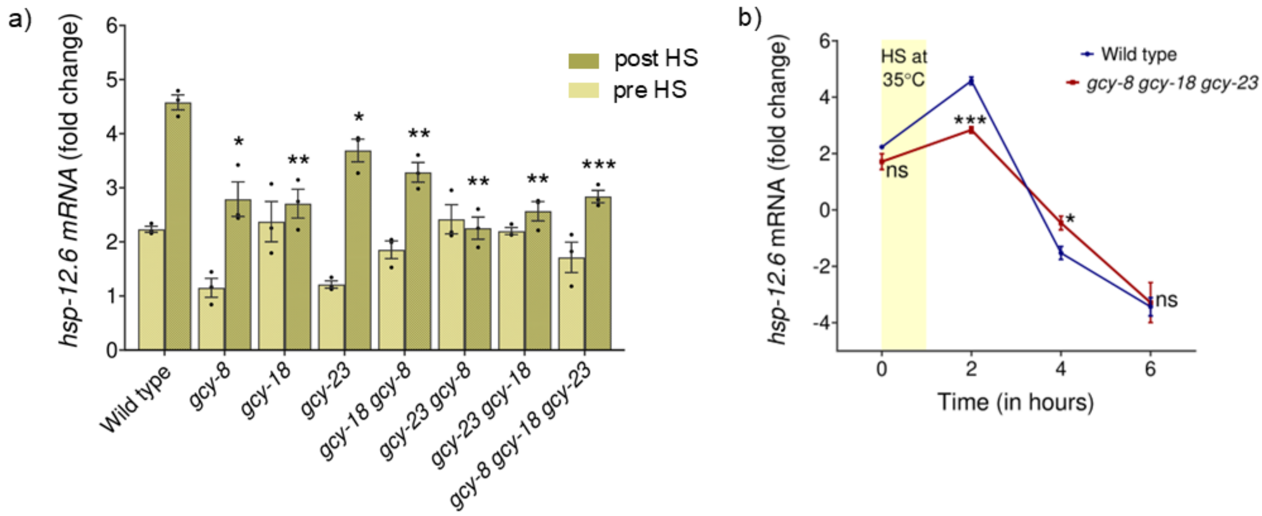


Figure S1. **AFD-rGCs regulate the expression and dynamics of *hsp-12.6* following thermal stress. (a)** Relative transcript levels of *hsp-12.6* in wild-type and across *gcy* single, double and triple mutants pre and post heat shock (35° C for 1 h). Expression levels were measured in worms collected 1 h post heat shock and comparisons are performed to post-heat shock levels in wild-type. **(b)** Time-course analysis of *hsp-12.6* mRNA levels in wild-type and *gcy-8 gcy-18 gcy-23* triple mutants at pre and post-heat shock (35°C for 1 h) time points (0, 2, 4 and 6 h). The shaded yellow region represents the heat shock period. All values represent mean ± SEM from at least three biological replicates. Statistical significance was determined using Welch's t-test comparing mutants to wild-type for each time point. ***$p < 0.001$, **$p < 0.01$, *$p < 0.05$, ns, not significant.

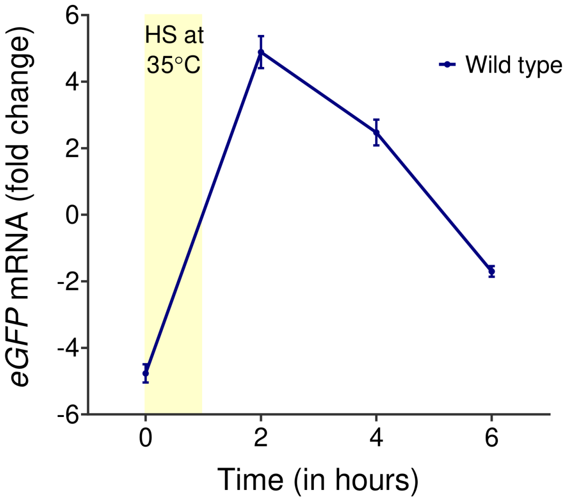


Figure S2. **Temporal dynamics of heat-induced *eGFP* mRNA expression in wild-type worms (TJ3001[zSi3001]).** Fold change in *eGFP* mRNA levels in wild-type animals before heat shock (0 h time point) and at time points 1 h post heat stress at 35°C (2, 4 and 6 h). The yellow shaded region indicates the heat shock period. Data represent mean ± SEM from at least three biological replicates.

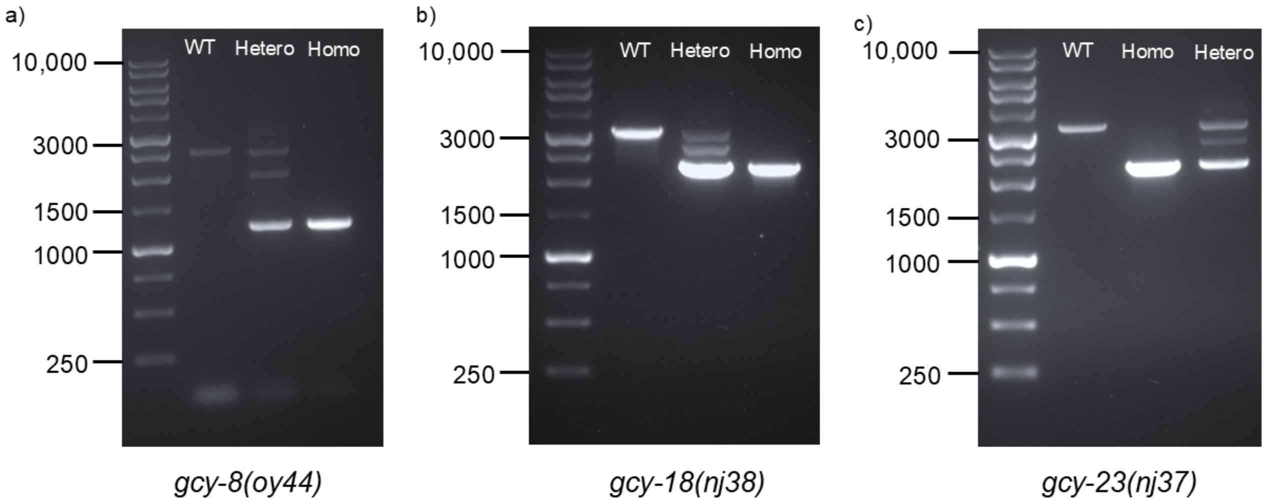


Figure S3. **Genotyping of *gcy* mutants crossed with strain TJ3001 to introduce *hsp-16.2p::GFP*.** Representative gel images showing single worm PCR-based genotyping assessment of wild-type (WT), heterozygous (Hetero) and homozygous (Homo) alleles for **a)** *gcy-8(oy44)*, **b)** *gcy-18(nj38)* and **c)** *gcy-23(nj37)*. DNA ladders with corresponding base pair sizes are shown on the left. Distinct band patterns were used to distinguish WT, heterozygous and homozygous animals for each allele.

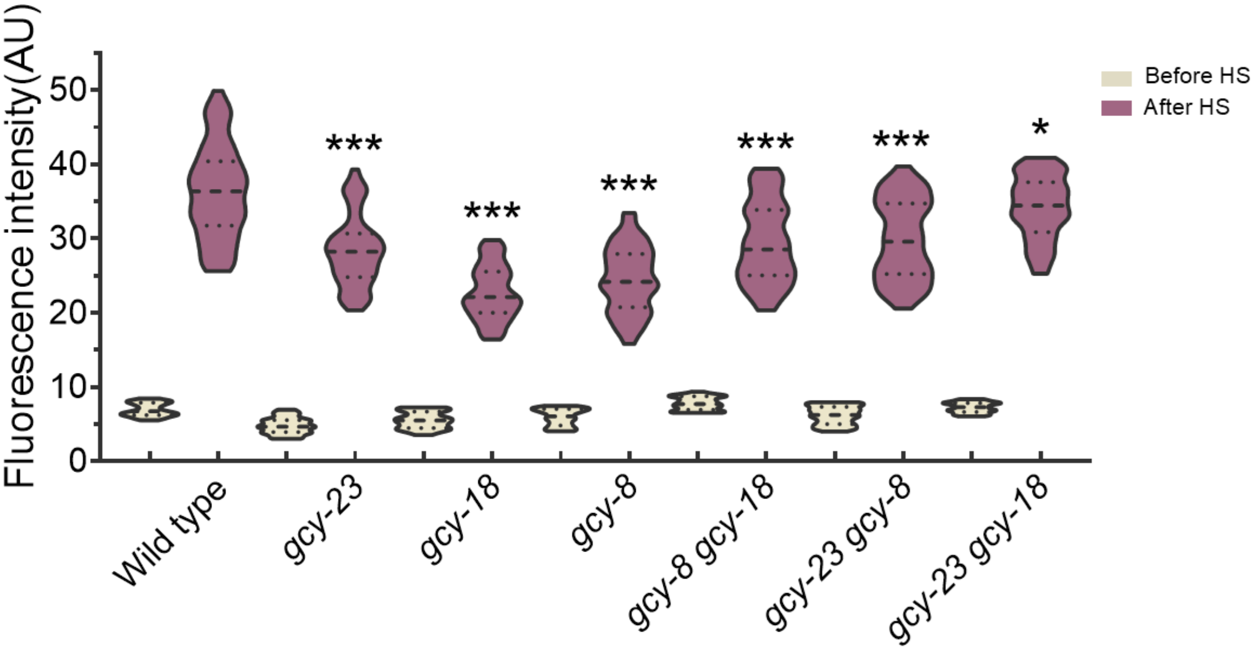


Figure S4. **HSP-16.2 expression analysis by quantification of GFP fluorescence intensity in AFD-rGC mutants before and after heat shock.** Fluorescence intensity of GFP in wild-type and *gcy* single and double mutant animals before and after heat shock at 35°C for 1 h. Each experiment was repeated at least three times with 30–40 animals imaged per condition. Fluorescence intensity is represented in arbitrary units (AU). Statistical comparisons were made using Student's t-test between each mutant and wild-type post heat shock (*p $<$ 0.05, ***p $<$ 0.001).

| STRAINS | SOURCE | IDENTIFIER |
|---|---|---|
| *C. elegans* strains | | |
| Wild type, Bristol | CGC | N2 |
| *gcy-8(oy44) IV* | CGC | IK800 |
| *gcy-18(nj38) IV* | CGC | IK429 |
| *gcy-23(nj37)IV* | CGC | IK427 |
| *gcy-8(oy44) gcy-18(nj38) IV* | Mori Lab, Nagoya University | IK746 |
| *gcy-23(nj37) gcy-8(oy44) IV* | Mori Lab, Nagoya University | IK747 |
| *gcy-23(nj37) gcy-18(nj38) IV* | Mori Lab, Nagoya University | IK748 |
| *gcy-23(nj37) gcy-8(oy44) gcy-18(nj38) IV* | CGC | IK597 |
| *zSi3001 [hsp-16.2p::GFP::unc-54 + Cbr-unc-119(+)] II* | CGC | TJ3001 |
| *unc-119(ed3) III; oxTi581 [eft-3p::tdTomato::H2B:: unc-54 3'UTR + Cbr-unc-119(+)] IV* | CGC | EG7935 |
| *gcy-23(nj37) gcy-8(oy44) gcy-18(nj38) IV; zSi3001 [hsp-16.2p::GFP::unc-54 + Cbr-unc-119(+)] II* | This study | |
| *gcy-8(oy44) IV; zSi3001[hsp-16.2p::GFP::unc-54 + Cbr-unc-119(+)] II* | This study | |
| *gcy-18(nj38) IV; zSi3001[hsp-16.2p::GFP::unc-54 + Cbr-unc-119(+)] II* | This study | |
| *gcy-23(nj37) IV; zSi3001[hsp-16.2p::GFP::unc-54 + Cbr-unc-119(+)] II* | This study | |
| *gcy-8(oy44) gcy-18(nj38) IV; zSi3001[hsp-16.2p:: GFP::unc-54 + Cbr-unc-119(+)] II* | This study | |
| *gcy-23(nj37) gcy-8(oy44) IV; zSi3001[hsp-16.2p:: GFP::unc-54 + Cbr-unc-119(+)] II* | This study | |
| *gcy-23(nj37) gcy-18(nj38) IV; zSi3001[hsp-16.2p:: GFP::unc-54 + Cbr-unc-119(+)] II* | This study | |
| Bacterial strains | | |
| E. Coli OP50 | CGC | WB OP50 |

Table S1. List of *C. elegans* and bacterial strains used in this study along with their sources and strain identifiers.

| Target | Forward | Reverse |
|---|---|---|
| *cdc-42* | TCCACAGACCGACGTGTTTC | AGGCACCCATTTTTCTCGGA |
| *pmp-3* | GTTCCCGTGTTCATCACTCAT | ACACCGTCGAGAAGCTGTAGA |
| *ire-1* | TACTTGCCACCACGGAGACC | CGTTGCCATCGTCATCATTG |
| *hsp-16.2* | TCCATCTGAGTCTTCTGAGATTGTT | TGATAGCGTACGACCATCCAAA |
| *hsp-16.1* | ATGGCTCAGATGGAACGTCA | TGGCTTGAACTGCGAGACAT |
| *hsp-16.48* | CTCATGCTCCGTTCTCCATT | ACAATCTCTCCAATATTGTCGGA |
| *hsp-70* | TCGATGAAGTTGTCTTGGTTGG | AGGCTACTGCTTCGTCTGGATT |
| *hsp-12.6* | ATGATGAGCGTTCCAGTGATG | TGATCATCGTCGTCGAGGAC |
| *hsp-90* | ATTCGCTACCAGGCACTCAC | TGGTAAGGGTCTTTTCCTCCT |
| *gcy-8* | AATCCCAAAGAAGCTGGCCTAC | GACAGCTAGTACATGGGTGAGC |
| *gcy-18* | TCGAAGAAATCATGCACAGG | CGTAGGCTGGTAGGAGTTGG |
| *gcy-23* | ATAGGCAATAACGAGGTGCG | AACTGGATTCTGGCCGTCTA |
| *egfp* | CCTGTCCACACAATCTGCCC | TGGTCTCTCTTTTCGTTGGGAT |

Table S2. List of gene-specific primers used in this study.

| REAGENT or RESOURCE | SOURCE | CATALOG DETAILS |
|---|---|---|
| Chemicals | | |
| RNAiso Plus (Trizol) | Takara | Cat# 9109 |
| Chloroform | Sigma | Cat# 288306 |
| Isopropyl alcohol | Sigma | Cat# I9516 |
| Glycogen | Thermo Fisher Scientific | Cat# R0551 |
| TE buffer (pH 8.0) | Thermo Fisher Scientific | Cat# AM9849 |
| Ethanol (absolute) | LABNOL | Cat# 5268-39 |
| RNA Loading Dye, (2X) | NEB | Cat# B0363S |
| KCl | Himedia | Cat# MB043 |
| MgCl2 | Himedia | Cat# MB040 |
| Tris (pH 8.3) | Himedia | Cat# ML156 |
| NP-40 | Himedia | Cat# MB143 |
| Tween-20 | Himedia | Cat# MB067 |
| Gelatin | Himedia | Cat# GRM019 |
| Proteinase K | NEB | Cat# P8107S |
| 50X TAE Buffer | Himedia | Cat# ML016 |
| LongAmp Taq DNA Polymerase | NEB | Cat# M0323S |
| LongAmp Taq DNA Buffer | NEB | Cat# B0323S |
| Deoxynucleotide (dNTP) Solution Mix | NEB | Cat# N0447S |
| 1Kb DNA Ladder RTU | GeneDireX | DM010-R500 |
| Critical commercial assays | | |
| RapidOut DNA Removal Kit | Thermo Fisher Scientific | Cat# K2981 |
| iScript™ cDNA Synthesis Kit | BioRad | Cat# 1708891 |
| iTaq™ Universal SYBR Green Supermix | BioRad | Cat# 1725124 |
| Softwares | | |
| FIJI | Schindelin et al., 2012 [1] | https://imagej.net/Fiji#Downloads |
| SnapGene | - | https://www.snapgene.com/ |
| OASIS 2 | Han et al., 2016 [2] | https://sbi.postech.ac.kr/oasis2/ |

Table S3. Key resources and reagents used in this study.

[1] Johannes Schindelin, Ignacio Arganda-Carreras, Erwin Frise, Verena Kaynig, Mark Longair, Tobias Pietzsch, Stephan Preibisch, Curtis Rueden, Stephan Saalfeld, Benjamin Schmid, Jean-Yves Tinevez, Daniel James White, Volker Hartenstein, Kevin Eliceiri, Pavel Tomancak, and Albert Cardona. Fiji: an open-source platform for biological-image analysis. *Nat. Methods*, 9 (7):676–682, June 2012.

[2] Seong Kyu Han, Dongyeop Lee, Heetak Lee, Donghyo Kim, Heehwa G Son, Jae-Seong Yang, Seung-Jae V Lee, and Sanguk Kim. OASIS 2: online application for survival analysis 2 with features for the analysis of maximal lifespan and healthspan in aging research. *Oncotarget*, 7 (35):56147–56152, August 2016.
